# A unique single nucleotide polymorphism in Agouti Signalling Protein (*ASIP*) gene changes coat colour of Sri Lankan Leopard (*Panthera pardus kotiya*) to dark black

**DOI:** 10.1101/2022.06.02.494537

**Authors:** Meegasthanne Gamaralalage Chandana Sooriyabandara, Asitha Udaya Bandaranayake, Shyaman Jayasundara, Hathurusinghe Arachchilage Bhagya Madhushani Hathurusinghe, Marasinghe Sumanasirige Leslie Ranjan Pushpakumara. Marasighe, Gajadeera Arachchige Tharaka Prasad, Vithana Pathirannehalage Malaka Kasun Abeywardana, Manoj Akalanka Pinidiya, Rajapakse Mudiyanselage Renuka Nilanthi, Pradeepa Chandani Gunathilake Bandaranayake

**Affiliations:** Department of Wildlife Conservation, 811/A, Jayanthipura Road, Battaramulla, Sri Lanka; Department of Computer Engineering, Faculty of Engineering, University of Peradeniya, Peradeniya, Sri Lanka; Agricultural Biotechnology Centre, Faculty of Agriculture, University of Peradeniya, Peradeniya, Sri Lanka

**Keywords:** - black panther, comparative genomics, coat colour, Sri Lankan leopard

## Abstract

The Sri Lankan Leopard (*Panthera pardus kotiya*) is an endangered subspecies restricted to isolated and fragmented populations in Sri Lanka. Among them, the melanistic leopards have been recorded on rare occasions. The existing literature suggests that melanism evolved several times in the Felidae family, with three separate species revealing distinct mutations. Nevertheless, the mutations in the remaining species, including Sri Lankan black leopard, are unknown. We used reference-based assembled the nuclear genomes of Sri Lankan normal and black leopard and *de novo* assembled mitogenomes of the same to investigate the genetic basis, adaptive significance, and evolutionary history of the Sri Lankan black leopard. Our data suggested coalescence time of Sri Lankan regular and black leopards at ∼0.5 Million years, sisters to *Panthera pardus* lineage. Interestingly, in the black leopard, a single nucleotide polymorphism in exon-4 possibly completely ablates Agouti Signaling Protein (ASIP) function. Existing genomic data suggest new a species-specific mutation of the ASIP gene in the Felidae family, contributing to naturally occurring colouration polymorphism. As such, the Sri Lankan black leopard and normal leopard probably evolved from the same ancestor, while the mutation in the ASIP gene resulted in black coat colour. This rare mutation could be adaptable to the environment that back leopards reported, camouflage, with a likelihood of recurrence and transmission to future generations. However, protecting this sensitive environment is critical for the conservation of the existing populations and providing breeding grounds.

## BACKGROUND

Leopards are considered critically endangered in many habitats (1), including Sri Lanka (2). *Panthera pardus kotiya,* the Sri Lankan leopard, is one of nine unique sub-species and the second remaining island leopard in the world (1, 3). Leopard subspecies are recognized based on genetic, morphological, and geographical information (4). Based on some DNA analysis, the nine subspecies recognized in 1996 are; *Panthera pardus pardus* (Linnaeus, 1758): Africa, *Panthera pardus nimr* (Hemprich & Ehrenberg, 1833): Arabia, *Panthera pardus saxicolour* (Pocock, 1927): Central Asia, *Panthera pardus melas* (Cuvier, 1809): Java, *Panthera pardus fusca* (Meyer, 1794): Indian sub- continent, *Panthera pardus delacourii* (Pocock, 1930): southeast Asia into southern China, *Panthera pardus japonensis* (Gray, 1862): northern China, *Panthera pardus orientalis* (Schlegel, 1857): Russian Far East, the Korean peninsula and north-eastern China and *Panthera pardus kotiya* (Deraniyagala, 1956): Sri Lanka, this subspecies is endemic to the island (1, 5). In addition, the leopard populations consist of significant genetic and morphological variation across, and in many cases, genetic patterns do not correspond to geographical variation recorded for the particular subspecies (1,6–10). Among them, melanistic leopard forms occur throughout its range, mostly in humid areas (11).

Reconstruction of the phylogenetic history of the leopards provides information on species, subspecies, and population genetic status, essential for the conservation of these threatened animals (12). The unique features of the leopards, for example, rosette pattern, spot size, and coat assist in resolving phylogeography relationships between the leopards (13–15). Molecular data have strengthened the conservation studies on other cat species; snow leopards, lions, and tigers (16, 17). Similarly, genetic analyses provide taxonomic guidelines for subspecies determination in leopards (1). Previous studies included morphological traits and three molecular genetic methods allozymes (3), mitochondrial DNA (18) restricted fragment length polymorphism (mtDNA–RFLP), and minisatellites to resolve six phylogeographic groups of leopards, including *P.panthera kotiya* (1). Intraspecific genetic variation of West Asian leopards was analyzed using the mitochondrial gene, NADH dehydrogenase subunit 5 (*NADH-5*), which exhibits a relatively high rate of mutation in leopards (19). Mitochondrial markers have been employed to identify leopard coalescence and lineage sorting processes (1,12,18,20). The MtDNA has been used to address questions of genetic diversity, phylogeography, and population evolution within Asian leopards, especially the Sri Lankan leopard (21–24).

The Sri Lankan leopard has isolated and evolved from dominant intra-guild competition at least since Sri Lanka split off from the Indian subcontinent ∼5,000–10,000 ybp (25–27). It is also the largest land carnivore and apex predator on the island (2,3,28,29). The Sri Lankan leopard range encompasses a variety of habitats, including the montane, sub-montane, tropical rain, monsoonal evergreen, and arid zone scrub forests (2,2,29,29–31). Despite its widespread distribution on the island, habitat loss, forest fragmentation, trapping, and hunting pose rising threats to the vulnerable sub-species (27,32,33).

Other than the regular coat colour leopards, a countable number of black leopards have also been reported from various parts of the island. Apart from Sri Lanka, black leopards have been reported in South-western China, Burma, Assam, and Nepal; from Travancore and other parts of Southern India, and are also common in Java and the Southern part of Malay. They are less common in tropical Africa but reported from Ethiopia (formerly Abyssinia), the forests of Mount Kenya, and the Aberdares (34, 35). The sightings have been rare due to the low population and the animal’s solitary nature. In Sri Lanka, black leopards have reliably been reported from Hiniduma, Sinharaja Forest, Warakadeniya, and moist densely-forested areas of hill country (36). During the last decade, two black leopards were recorded, both were victims of crucial deaths due to traps (36). Some environmentalists claim that the black leopard died in 2020 after injuries was the last in Sri Lanka.

The density of melanin and the distribution of melanin types on individual hairs determine the coat colour in mammals (37). Extreme phenotypic changes in the coat colour patterns have been reported in thirteen Felidae species. So far, only four different melanistic Felidae species have been analyzed. Those exhibit unique mutations associated with two genes; Melanocortin-1 receptor (*MC1R*) (35, 38) and Agouti Signalling Protein (*ASIP*) (35,39,40). In leopards, melanism is caused by a recessively inherited mutation in the *ASIP* gene, which leads to a non-sense mutation ablating *ASIP* function and thus induces black pigmentation (40). However, in jaguar, it has been inherited by a dominant trait caused by a 15-base pair deletion in the *MC1R* gene that leads to a gain of function mutation favoring the production of eumelanin (41). A SNP has been identified at exon 4 of *ASIP* gene in *Pardofelis temminckii* at position 384 resulting in a non-synonymous substitution associated with black coat colour (40). Nevertheless, mutations related to melanism of other leopard subspecies are yet to be studied. A broader assessment of its evolutionary history and adaptive significance assists strong conservation strategies. For example, the phylogeography of the Sri Lankan black leopard, whether it is a subspecies or a mutant of the Sri Lankan leopard, *P. p. kotiya*, is a serious discussion among environmental groups.

Therefore, we utilized data obtained from the dead black leopard in 2020, a regular *P. p. kotiya,* and published data to assess genetic variation and genetic differentiation in contemporary leopard populations. Here, we assembled the complete reference guide genome and *de novo* mitogenome of *P. p. kotiya* (regular coat colour) and black coat colour to study its structure and functional components for example subunit 5 of the *NADH* gene, and to compare with members of the subspecies *P.pardus.* Our study aimed to utilize the molecular data, for a better understanding genetic structure of the leopards in Sri Lanka, identify the possibility of sub speciation present within the island, establish baseline information for future monitoring, and identify genes and mutations for functional melanism. Scientific knowledge on melanism and species assessment in Sri Lanka help to prioritize the conservation efforts of leopards in the Island and around the globe.

## MATERIALS AND METHODS

### Ethical clearance

The study was conducted under the research permit WL/3/2/2017/1, issued by the Research Committee of the Department of Wildlife Conservation, Sri Lanka (DWC). We followed relevant guidelines and regulations for animal research approved by the committee. The DNA samples for sequencing were exported under the Convention on International Trade in Endangered Species of Wild Fauna and Flora (CITES) permit obtained from the DWC.

### Field Observations and Morphology Analysis

Ten morphometric measurements were taken to the nearest 0.1 cm from a dead black leopard, died on 29^th^ May 2020 after recovering from a trap at Hatton in Nuwara Eliya district (N- 6.8304983, E- 80.532660). The same animal has been reported in the cameras on 10^th^ December 2019, set up by the DWC. The regular leopard was found at Yala National Park (GPS: N- 6.308977, E- 81.423837. A non-stretch tape was used to measure body, tail, and head to tail length and a graduated wooden sliding calliper to record shoulder height and tooth dimensions. A photograph of the right-side profile of each leopard was taken for identification.

### Sample collection and DNA extraction

About 5 ml of blood was obtained during the post-mortem, from the heart blood to a tube containing EDTA using a sterile syringe. A similar amount of blood was drawn from a captive wild leopard from safeness vein at capture for medical treatment. The DNA was extracted using the Wizard Genomic DNA Purification kit (Promega, A1120) following the manufacturers’ guidelines and quantified using the Nanodrop 2000 Spectrophotometer (Thermo Scientific). The quality was assessed with Nanodrop readings and running on a 1% agarose gel. Samples were shipped to Admera Health Biopharma Services, New Jersey, USA for sequencing.

### Genomic library preparation and sequencing

At the sequencing facility, the genomic DNA was quantified with Qubit 2.0 DNA HS Assay (ThermoFisher, Massachusetts, USA) and assessed the quality with Tapestation genomic DNA Assay (Agilent Technologies, California, USA). The sequencing libraries were prepared using the KAPA Hyper Prep kit (Roche, Basel, Switzerland) per the manufacturer’s recommendations with Illumina® 8-nt dual-indices. The quantity of the final libraries was assessed by Qubit 2.0 (ThermoFisher, Massachusetts, USA), and quality was assessed by TapeStation D1000 ScreenTape (Agilent Technologies Inc., California, USA). Libraries were sequenced on an Illumina HiSeq (Illumina, California, USA) with a read length configuration of 150 PE for 600 M PE reads per sample (300 M in each direction). The preliminary quality assessment of the whole genome sequencing libraries was performed with FastQC v0.11.8(68). The FastQC reports were then concatenated using MultiQC v1.9 (69) (36) to create a single diagnostic report containing GC content, sequence quality, duplication level, and adapter content.

### *De novo* assembly and annotation of *P. p. kotiya* and black leopard

The *P. p. kotiya* and black leopard mitogenomes were assembled and annotated using the Mitoz v2.3 program. We extracted 5GB of reads from whole genome datasets using the BBMap reformat.sh tool, as recommended by the tool. The generated mitochondrial genomes and annotations were compared with the already available *P. pardus* mitogenomes (Table 1). The assembled genomes were visualized with OrganellarGenomeDRAW v1.3.1 (72) software.

**Table 1:**
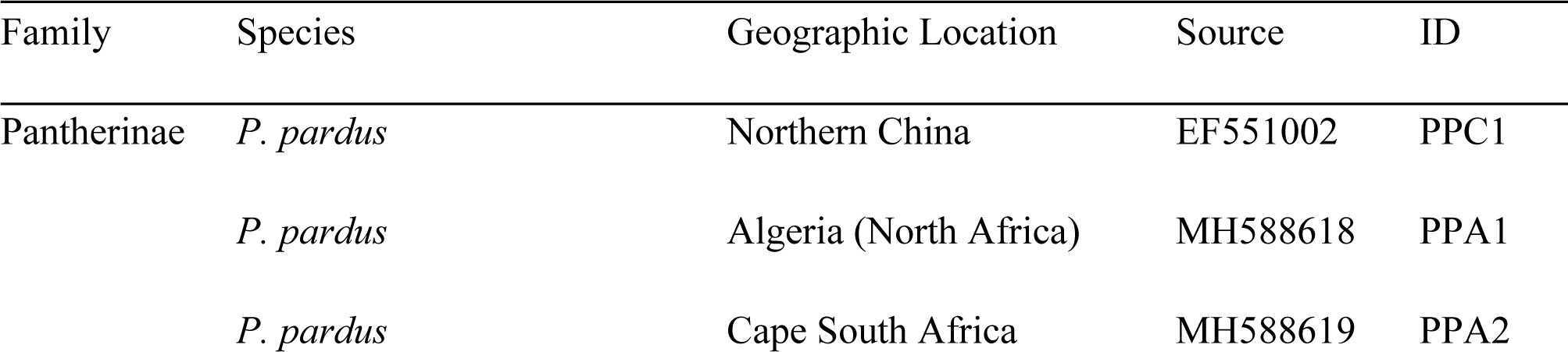
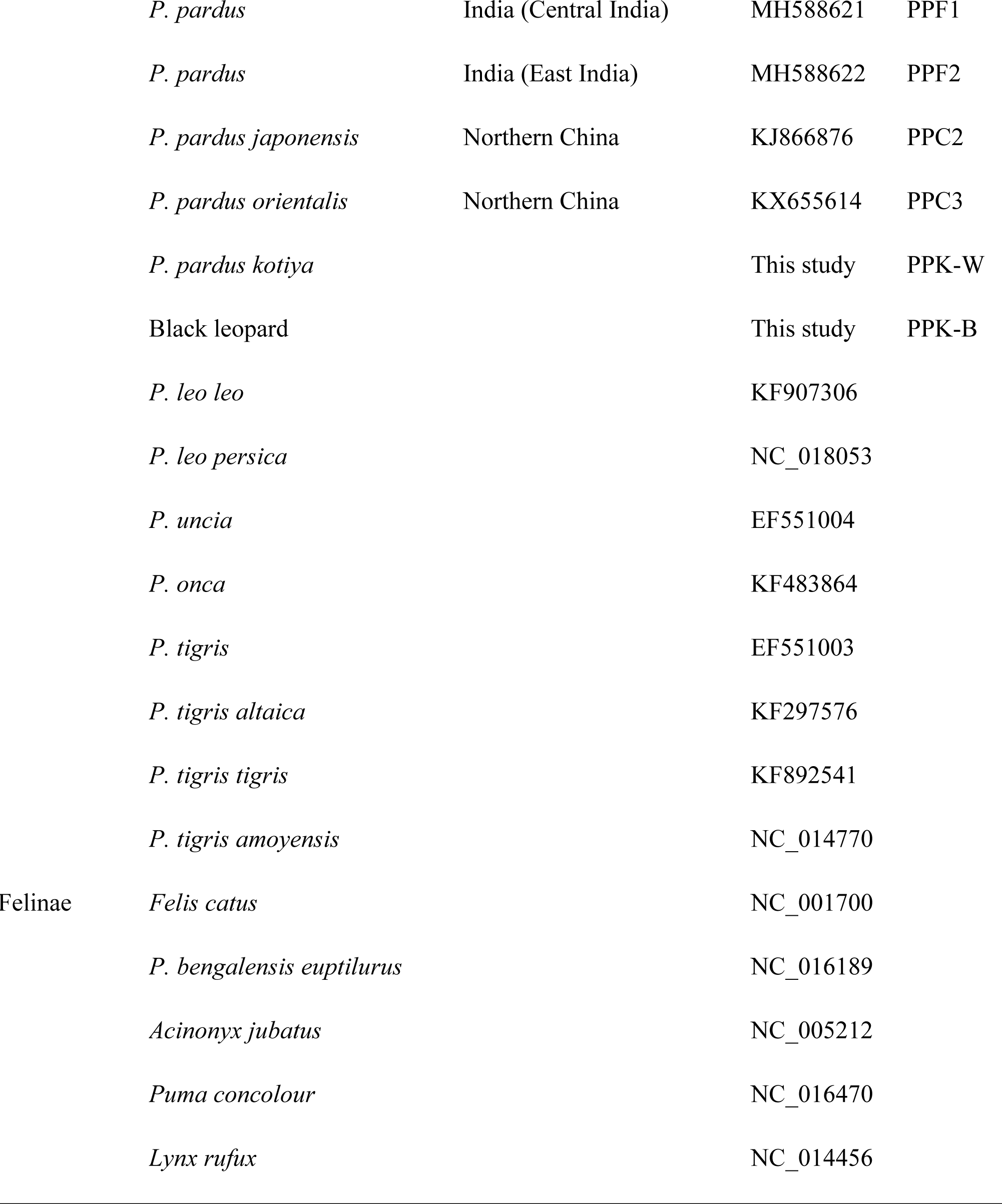
Pantherinae and Felinae mitochondrial genomes included

| Family | Species | Geographic Location | Source | ID |
| --- | --- | --- | --- | --- |
| Pantherinae | *P. pardus* | Northern China | EF551002 | PPC1 |
|  | *P. pardus* | Algeria (North Africa) | MH588618 | PPA1 |
|  | *P. pardus* | Cape South Africa | MH588619 | PPA2 |
|  | *P. pardus* | India (Central India) | MH588621 | PPF1 |
|  | *P. pardus* | India (East India) | MH588622 | PPF2 |
|  | *P. pardus japonensis* | Northern China | KJ866876 | PPC2 |
|  | *P. pardus orientalis* | Northern China | KX655614 | PPC3 |
|  | *P. pardus kotiya* |  | This study | PPK-W |
|  | Black leopard |  | This study | PPK-B |
|  | *P. leo leo* |  | KF907306 |  |
|  | *P. leo persica* |  | NC_018053 |  |
|  | *P. uncia* |  | EF551004 |  |
|  | *P. onca* |  | KF483864 |  |
|  | *P. tigris* |  | EF551003 |  |
|  | *P. tigris altaica* |  | KF297576 |  |
|  | *P. tigris tigris* |  | KF892541 |  |
|  | *P. tigris amoyensis* |  | NC_014770 |  |
| Felinae | *Felis catus* |  | NC_001700 |  |
|  | *P. bengalensis euptilurus* |  | NC_016189 |  |
|  | *Acinonyx jubatus* |  | NC_005212 |  |
|  | *Puma concolour* |  | NC_016470 |  |
|  | *Lynx rufus* |  | NC_014456 |  |

### MtDNA divergence dating and Nuclear DNA phylogeny

We downloaded 15 Pantherinae mitogenomes and 5 Felinae mitogenomes (Table 1). However, we did not include the partial mitogenomes (MH588618-MH588622) in the divergence dating analysis. The mtDNA D-loop region was excluded from the alignment due to its complexity.We used MAFFT (v.7.450) (73) with its default configuration for multiple sequence alignment. The Bayesian evolutionary analysis was carried out using the BEAST v2.6.5 (74) software package with Markov Chain Monte Carlo (MCMC) procedure to estimate posterior distributions of divergence times and evolutionary dates. The HKY model of evolution and gamma+invariant rate heterogeneity models with a relaxed lognormal clock were used to calculate the substitution rate, while the Yule node calibration technique was used to calculate the divergence. The posterior distributions were calculated using Markov chain Monte Carlo (MCMC) sampling with 100,000,000 steps and 10,000 step logging. We imposed monophyletic constraints for the node to calibrate evolutionary rates, and the times to the most recent common ancestor (TMRCA) of Pantherinae and Felinae with normal priors (10.78 ± 1.87 Mya) (75) were used for the node calibration. Tracer v1.7.2 (76) was used to examine the BEAST output. After discarding 10% MCMC steps as postburn-in, the maximum credibility tree was obtained using TreeAnnotator v2.6.4 (from the BEAST package) and visualized with FigTree v1.4.4 (77).

### *NADH-5* gene phylogeny

The nucleotide variation of the *NADH-5* gene of the Sri Lankan leopard samples was compared to that of the Indian leopards. In addition to NADH-5 orthologs retrieved from the PPK-W and PPK-B mitogenomes, we included NADH-5 gene sequences of Sri Lankan and Indian leopards available in the GenBank database (Table 2). A whole-genome sequencing dataset (ERR5671301) of a Sri Lankan leopard was also downloaded from the National Center for Biotechnology Information Sequence Read Archive (NCBI SRA), and reads were mapped to the *NADH-5* sequence (AY035267) of *P. pardus kotiya* using the BWA MEM (v0.7.17) aligner (78). The Geneious Prime 2020.1.2 program was used to generate the consensus sequence, that was included in the analysis. The *NADH- 5* orthologs of *P. tigris* (EF551003) and *P. leo* (KF907306) were identified as the outgroup (Table 1). We used the MEGA X v10.2.4 (79) software to identify the best model of nucleotide substitution based on the Bayesian Information Criterion (BIC). The Kishino and Yano (HKY) + G model of nucleotide substitution (gamma distribution with five rate categories) was identified as the most suitable. The maximum likelihood (ML) tree for *the NADH-5* gene was generated using MEGA X, setting the bootstrap to 1,000 replicates. Additionally, a Bayesian analysis was conducted using Mr Bayes v3.2.6 (80), using the HKG85 substitution model with a chain length of 1,100,000 and 100,000 burn-in. PopArt v1.7 (81) was used to construct a Median Joining haplotype network (ε =0) of all the sequences from Asian leopards.

**Table 2.**
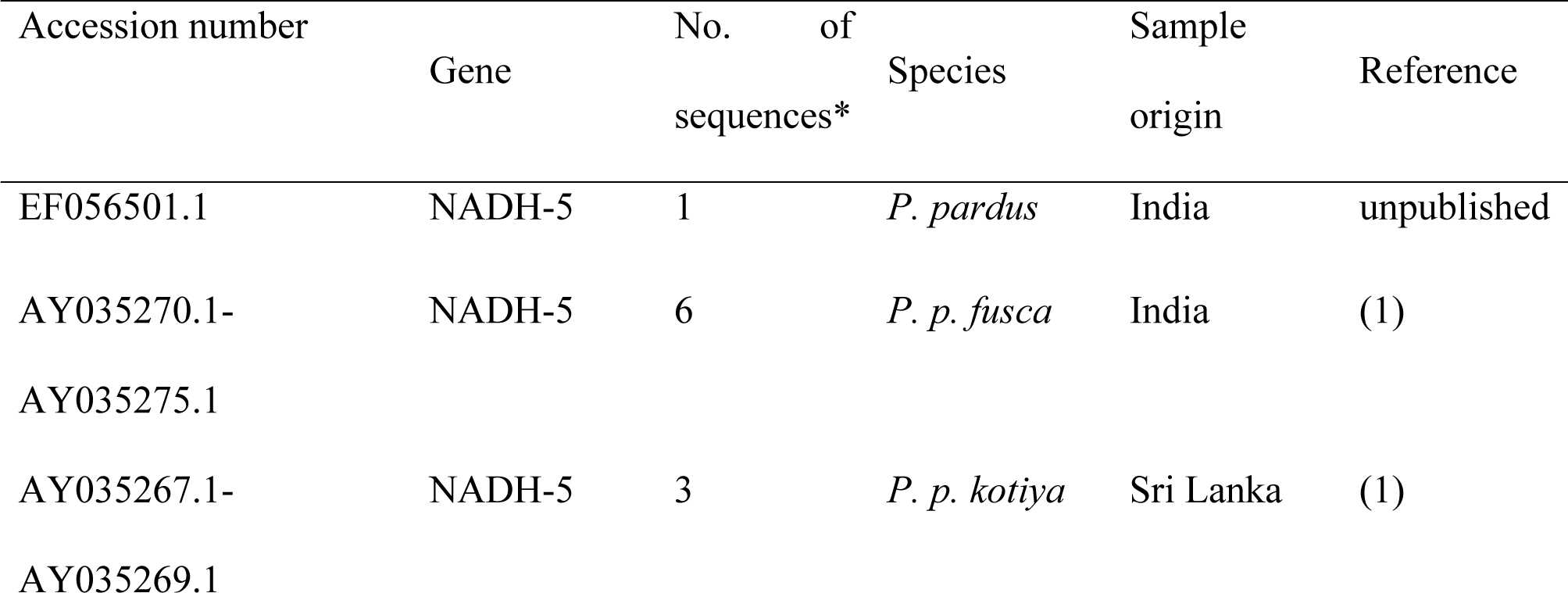
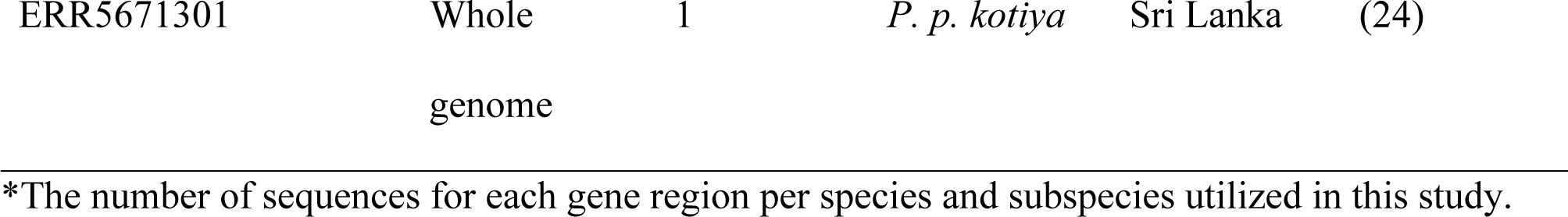
Leopard sequences obtained from Genbank.

**Table 3.** Nucleotide distance between Panthera pardus mitogenomes (excluding the D-Loop region)

| Mitogenome ID | PPA1 | PPA2 | PPF1 | PPF2 | PPC1 | PPC2 | PPC3 | PPK-W | PPK-B |
| --- | --- | --- | --- | --- | --- | --- | --- | --- | --- |
| PPA1 (North Africa) |  | 129 | 463 | 432 | 493 | 456 | 536 | 442 | 443 |
| PPA2 (South Africa) | 129 |  | 522 | 411 | 474 | 439 | 526 | 406 | 407 |
| PPF1 (Central India) | 463 | 522 |  | 127 | 249 | 214 | 307 | 220 | 220 |
| PPF2 (East India) | 432 | 411 | 127 |  | 163 | 116 | 226 | 99 | 99 |
| PPC1 (Northern China) | 493 | 474 | 249 | 163 |  | 184 | 286 | 185 | 185 |
| PPC2 (Northern China) | 456 | 439 | 214 | 116 | 184 |  | 134 | 133 | 133 |
| PPC3 (Northern China) | 536 | 526 | 307 | 226 | 286 | 134 |  | 239 | 239 |
| PPK-W (Sri Lanka) | 442 | 406 | 220 | 99 | 185 | 133 | 239 |  | 6 |
| PPK-B (Sri Lanka) | 443 | 407 | 220 | 99 | 185 | 133 | 239 | 6 |  |

The pairwise nucleotide distance was calculated using the multiple sequence alignment created for the divergence dating analysis of complete mitogenome excluding D-loop region.

**Table 4.** Statistics of reference-based genome assembly of P. p. kotiya and Black leopard

|  | *P. p. kotiya* (PPK-B) | Black leopard (PPK-W) |
| --- | --- | --- |
| Total read pairs | 1,236,550,993 | 312,991,352 |
| Unmapped reads (%) | 264,233,004(10%) | 4,085,418 (0.65%) |
| Duplicates (%) | 18.31 | 7.20 |
| Percentage mean coverage | 95.44 | 95.25 |
| Mean depth | 98.79 | 32.65 |
| Mean mapping quality | 55.79 | 57.41 |
| Number of variant sites | 7,977,753 | 14,939,356 |
| Variant sites after filtering (%) | 4,385,747 (54.97%) | 43,86,959 (29.37%) |
| SNPs | 3,181,511 | 3,216,359 |
| MNPs | 184,201 | 162,838 |
| Indels | 973,127 | 957,130 |

### Reference-based assembly of *P. p. kotiya* genomes and identification of mutation

We used two whole genome sequencing (WGS) datasets for the reference-guided assembly with the chromosome-length *P. pardus* genome assembly (53, 54) published by the DNA Zoo team (an improved version of PanPar1.0 (82)) as the reference genome. The reads were mapped to the reference genome with the BWA MEM (v0.7.17) aligner (78), and Samtools’ fixmate (84) was used to correct the aligner’s flaws in read-pairing. Picards’ MarkDuplicates (85) was used to identify duplicate reads in the BAM file, and Samtools’ sort (84) was used to sort the mapped reads in positional order. We used Samtools’ coverage command to generate per scaffold statistics (i.e., mean depth and coverage). Freebayes v1.3.4 (86) was used for variant calling, and variants were filtered by a minimum quality, minimum depth, and maximum depth of 40, 5, and 50, respectively, using VCFtools v0.1.16 (87). The bcftools (v1.11) (88) stats command was used to generate genotype statistics, and the bcftools consensus command was used to generate the consensus genome sequence. We selected a set of nuclear genes that code for vital enzymes in melanin synthesis pathways. The objective was to identify whether genes in melanin synthesis pathway regulate and alter the melanin production. .As such, we retrieved Agouti Signaling Protein coding gene.(*ASIP)* (Gene ID 109272540), tyrosinase-related protein 1 coding gene (*TYRP1)* (Gene ID 109247101), Dopachrome Tautomerase Protein coding gene *DCT* (Gene ID 109264168), OCA2 melanosomal transmembrane protein coding gene.*OCA2* (Gene ID 109258721), Phospholipase C Beta 4 protein coding gene*PLCB4* (Gene ID 109274721), Premelanosome Protein coding gene, *PMEL* (Gene ID 109270307), and egl-9 family hypoxia inducible factor 1*EGLN1* (Gene ID 109248931) gene sequences of *P. pardus* from NCBI. We mapped the melanocortin 1 receptor gene *MC1R* gene (Gene ID 493917) of *Felis catus* to the *P. pardus* reference genome (83) (PanPar1.0 Ensembl version 104) using Bowtie2 aligner (89) to extract the *MC1R* ortholog of the *P. pardus*. The ortholog sequences of these genes were then identified and extracted from the two leopard genome assemblies using the BLASTN function. The reads of the Sri Lankan leopard WGS dataset (ERR5671301) from the NCBI were mapped to the coding DNA sequences using the bwa mem aligner (78), and their ortholog consensus sequences were generated using 50% strict calling in Geneious Prime 2020.1.2 software (http://www.geneious.com/). In addition to these sequences, we included the ortholog sequences of the considering genes, which are available in the NCBI. Finally, the multiple sequence alignment of the *ASIP*, *TYRP1*, and *MC1R* orthologs of four *Panthera* samples were executed using the MAFFT v.7.450 (79) with its default configuration. The protein structure of the mutated gene was predicted with RoseTTAFold (90).

## RESULTS

### Comparative morphology analysis of Regular coat colour and Black leopards

From visual data, Sri Lankan Leopard’s Spot and Rosette pattern was distinct from the other leopards (42). The shapes, sizes, and formations of spot and rosette formations are unique to every leopard (29, 43). We also compared the spot and rosette formation of animals included in the analysis. The morphological parameters of two animals included in the analysis were compared (Additional File 1: Table S1). Based on the analysis, the black leopard is a male animal of about 10 years old in 2020 while the regular leopard is a female animal of about 2 years in 2020. The regular coat colour leopard is striking rust yellow with blackish spots and rosettes. A darker orangish contrast between the spots create the rosettes in some areas of the coat. Rosettes were of different sized and shapes (Figure 1a). The black leopard also has a distinctive Rosette pattern visible even in the darker background (Figure 1b), hidden due to black pigmentation, were termed “ghost rosettes.”

**Figure 1:**
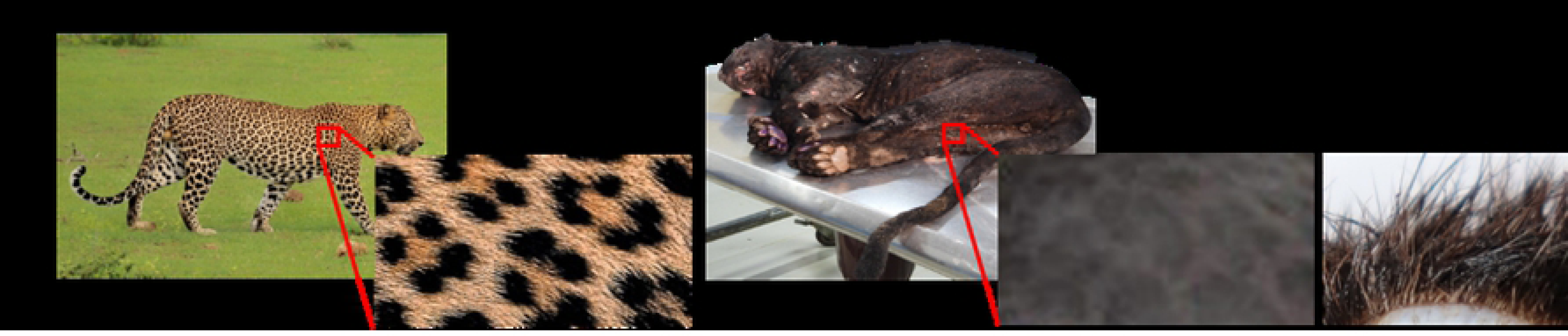
A visual Morphological differences between Sri Lankan regular and black leopards (a) The side view of Regular leopard (PPK-W) and the view of its spot and rosette pattern The side view of the Sri Lankan black leopard (PPK-B) and the view of its spot and rosette pattern. The spots appear to be still visible in the black background. The cross section of the skin of (PPK-B) with visible fur colouration

### Quality checking of the two whole genome sequencing datasets

We generated genome data for two leopard specimens excepting 100x genome coverage for black leopard and 30x genome coverage for regular coat *P. p. kotiya*. The obtained WGS datasets consist of 1236.6 and 313 million read pairs for *P. p. kotiya* and black leopard samples, respectively. These two datasets passed all the FastQC tests except the per sequence GC content test for the reverse reads of *P. p. kotiya* library (Additional File 2: Figure S1). Considering the good quality of the sequenced data, we used the original dataset for the downstream analysis.

### *De novo* mitogenome assembly and divergence dating

We *de novo* assembled the mitochondrial genome of a Sri Lankan leopard using the sequenced WGS dataset, and it resulted in a mitogenome of 16,847 bp long (Figure 2) with a 41% GC content. It encodes typical metazoan mtDNA genes, including 13 protein-coding genes (PCGs), 22 tRNA genes, two rRNA genes, and one non-coding control region (D-loop) (Additional File 3: Table S2). The gene order is identical to the *P. pardus* (EF551002) mitogenome. The total length of all protein-coding genes in the mitochondrial genome is 11,419bp (i.e., 67.8% of the assembled mitochondrial genome). The base composition is 31.8% of A, 26.4% of C, 14.6% of G, and 27.2% of T. The 16S rRNA and 12S rRNA of the rRNA gene are 1,577bp and 962bp, respectively. The length of 22 tRNA ranges from 58 bp (tRNA-S) to 75 bp (tRNA-L), while the control region of 1,424bp is located between tRNA-P and tRNA-F.

**Figure 2.**
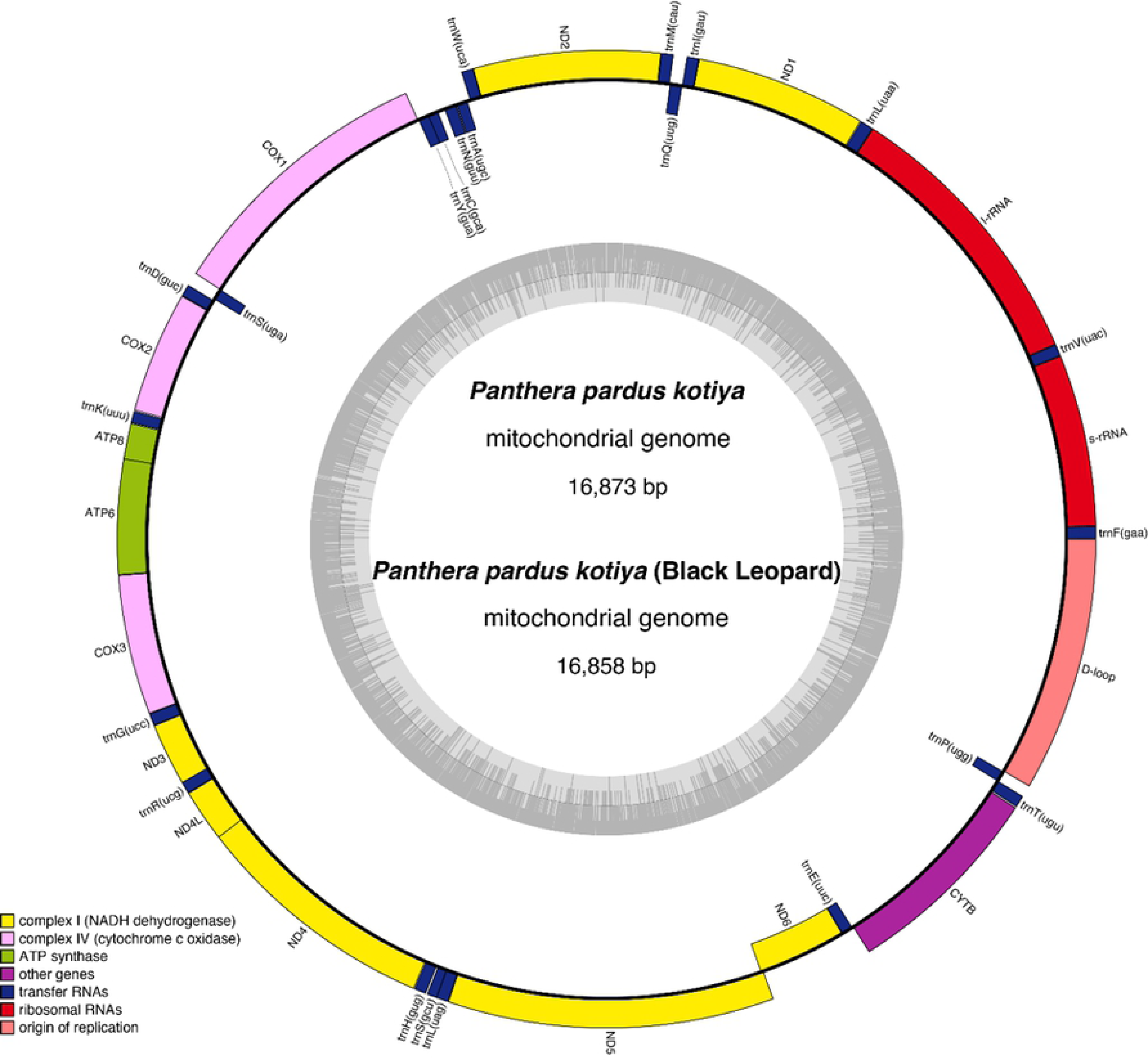
The mitogenome of *Panthera pardus kotiya* and black leopard. Gene components are colour-coded. The inner-circle graph depicts GC content across the genome.

The start and stop codons looked to be ubiquitous across all animals. Except for ND2, ND3, and ND5, which utilized ATC, ATA, and ATA, respectively, ATG was generally used as the start codon for all genes. TAA was commonly found as a stop codon while *ND2*, *ND3* ended with TAG, and *Cyt b* ended with AGA. The termination of the *Cyt b* gene with AGA is unique to mammalian taxa (44–46), as opposed to amphibians(47), reptiles(48, 49), Aves(50), and Nematoda(51). The stop codons in *ND4* and *COX3* are incomplete. These partial termination codons are ubiquitous in metazoan mitogenomes, and following transcription, poly A can convert them to full ones (TAA)(52).

Similarly, we assembled the mitochondrial genome of the black leopard (Figure 2). Together with previously published mitogenomes of Pantherinae and Felinae, we derived the phylogenetic relationships and estimated the coalescence times of the considered mitogenomes (Figure 3).

**Figure 3.**
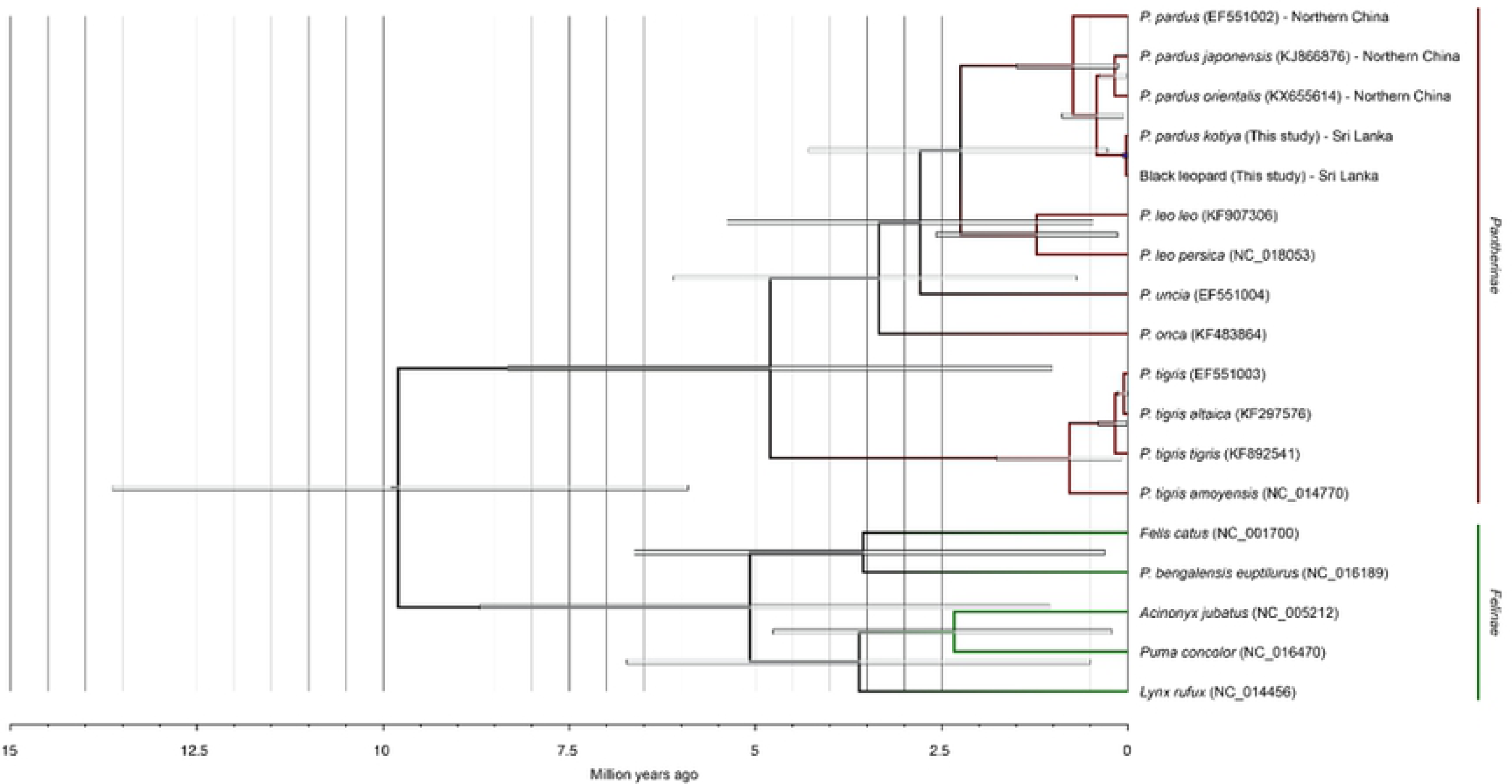
Mitochondrial phylogenetic tree of Pantherinae and Felinae. Node bars represent the 95% highest posterior density (HPD). Tree tips are labeled with the species name, accession number, and origin.

The Bayesian analyses based on 23 complete mitochondrial genomes, excluding the D-Loop control region, yielded the identical tree topology supported by a high bootstrap value (above 95%) and high posterior probabilities (above 0.95). The fossil-calibrated Bayesian analysis of the basal divergence time for all Felidae individuals at mitochondrial lineages at 10.78 Mya regenerates the history. BEAST analysis suggested divergence time of Felidae individual was approximately 9.62 Mya from the common ancestor. The divergence of *P. tigirs* was estimated approximately 4.9 Mya from the closely related Panthera species (ML, 95%; credibility interval CI 1.5-6.5 My). Divergence of other individuals; *P. leo* (lion) at ∼2.5 Mya (ML, 95%; CI 0.25-2.75 My), *P. uncia* (Snow leopard) at ∼2.0 Mya (ML, 95%; CI 0.9-5.5 My). and *P.onca* (Jaguar) *∼*3.5 Mya (ML, 95%; CI 1.1-6.5 My).

Estimation of divergence times of genus Panthera in the present study reconfirms earlier reports. The Markov Chain Monte Carlo (MCMC) analysis employed in BEAST revealed that *Panthera pardus* split from its sister species leopard at approximately 2.5 Mya. Our results support the deep bifurcation between Asian and African leopards, that has previously been proposed based on short mtDNA sequences. The divergence between the Asian leopards from the East-Central African subspecies is ∼1.25 Mya (ML, 95%; CI 0.25- 2.5My). The deepest roots within the *P.p. kotiya* (African origin) was estimated to be ∼0.5Mya (ML, 95%; CI 0.25- 1.1 My).

When the intraspecies divergence of the mitogenome of Africa, India, China, and Sri Lanka was examined, the two samples from Sri Lanka had the smallest nucleotide difference (Table ). When the D-Loop region is omitted, the *P. p. kotiya* and black leopard mitogenomes differ by 6 nucleotides, and five of these six SNPs were found in the *ND1*, *COX1*, *COX3*, and *CYTB* coding regions. Except for the SNP in the *ND1* coding region, the rest of the four mutations are synonymous. The two Indian leopards PPF1 and PPF2, are the second closest mitogenomes, with a base pair difference of only 127 nucleotides.

We used *NADH-5* sequences for the downstream analysis, because this region has a relatively high rate of mutation in leopards (18). When the sequences of 7 Indian leopards and 6 Sri Lankan leopards aligned, the ML analysis and Bayesian gene tree grouped the Sri Lankan and Indian leopards into separate groups (Figure 4). Despite the low bootstrap values in the ML tree, it is nearly identical to the Bayesian tree with greater posterior probabilities. Our PPK-W and PPK-B samples were clustered together with the Sri Lankan leopard samples. Furthermore, we focused on the links between our samples and the remaining samples in this group. A haplotype network analysis was performed on the same samples to determine genetic diversity between PPK-W and PPK-B samples and their relationship with the Sri Lankan leopards.

**Figure 4.**
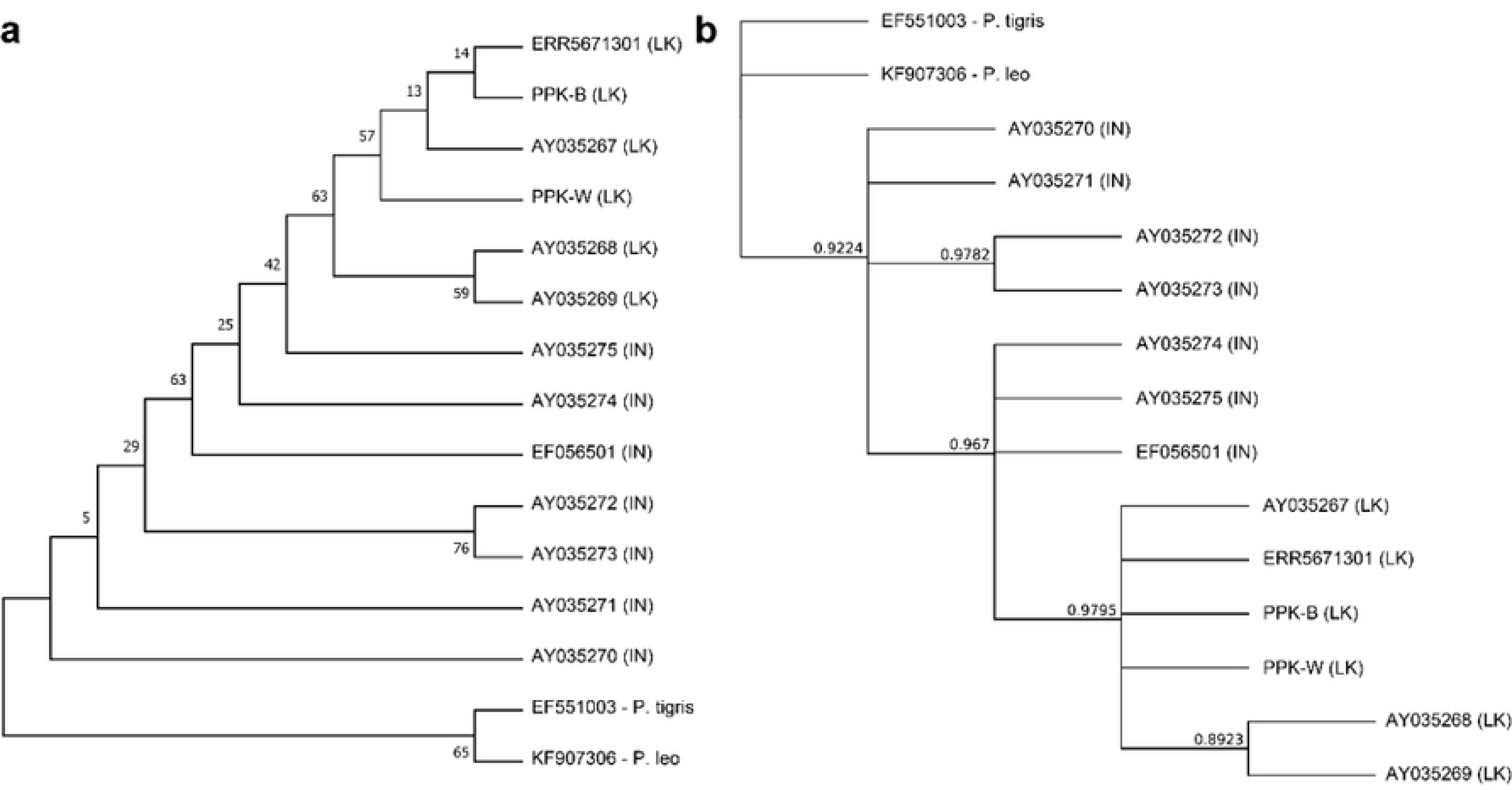
(a) Maximum likelihood (b) Bayesian *NADH-5* gene tree of 13 Asian leopards along with the *P. tigris* and *P. leo* outgroup. The bootstrap values and posterior probabilities are displayed next to the nodes, respectively. The origins of samples are given inside the brackets. LK – Sri Lanka, IN - India

In Sri Lankan leopard samples, the sequences provided here identified two unique haplotypes (Figure 5). This haplotype contains our two samples (PPK-W and PPK-B), as well as a previously reported whole-genome sequencing sample (ERR5671301) and was identical to the *Panthera pardus kotiya* haplotype (AY035267.1). Another haplotype was discovered with AY035268.1 and AY035269.1.

**Figure 5.**
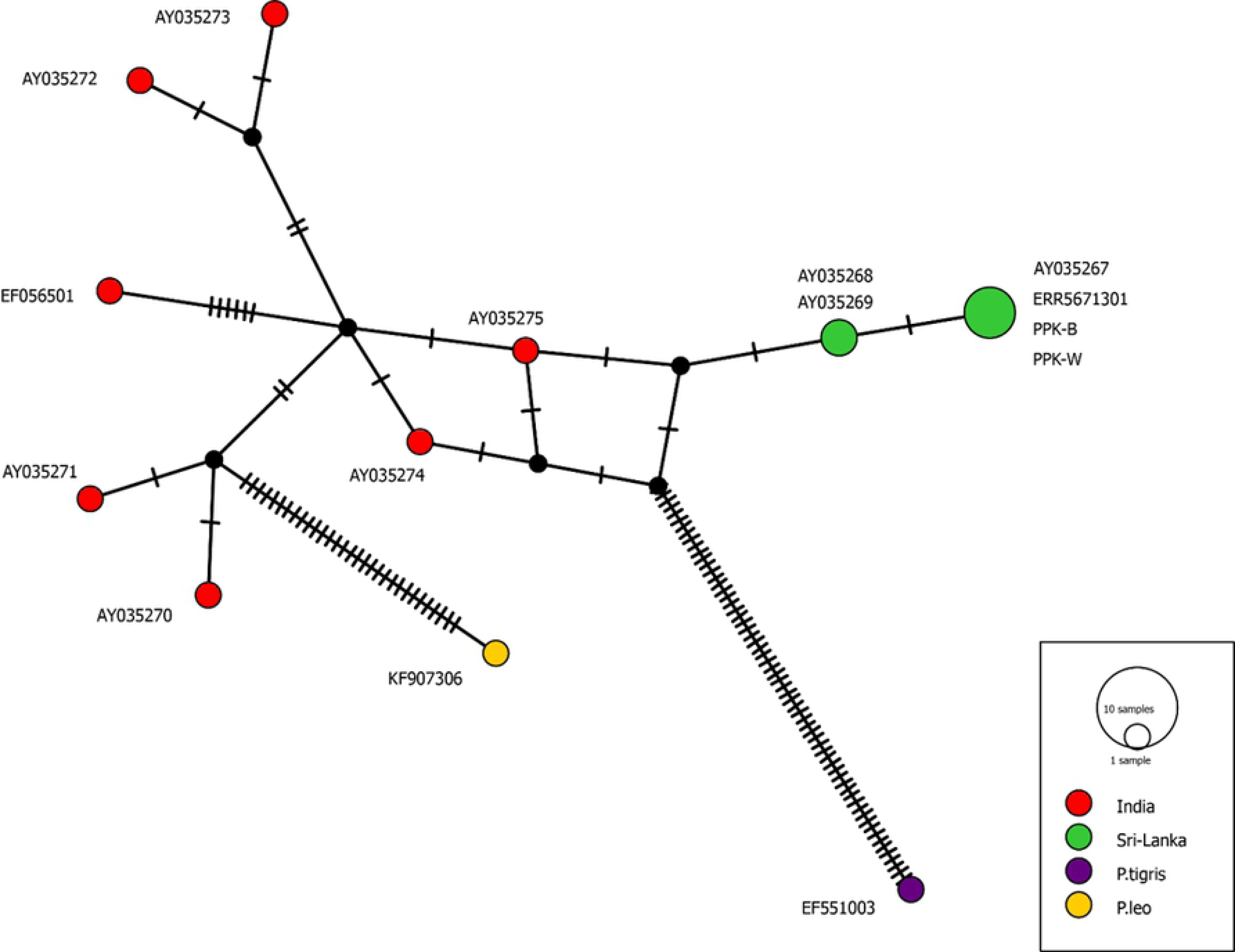
Median joining network analysis of the Asian leopards using *NADH-5* gene. The network includes 6 sequences of Sri Lankan leopards, and 7 sequences from Indian leopards. Each node represents a unique haplotype. Haplotypes are colour-coded based on the geographical location/species. Size of the circles represents the number of sequences with the same haplotype. Size of the node reflects the frequency of each haplotype.

### Reference guided assembly and Identification of *ASIP* mutations

We achieved 90% and 99% of mapping for PPK-B and PPK-W libraries, respectively (**Error! Reference source not found.**) for paired –end read mapping to the *P. pardus* reference genome (53, 54) The mean coverage and mean mapping quality were around 95% and 56, respectively for both the samples. As we expected, we have gained mean depths around 100x and 30x. After variant calling, we retrieved only the genotypes with a minimum of 5x depth. Since the sites with unusually high coverage can result in the incorrect mapping of reads from duplicated regions, we set the maximum depth to 50x and 100x for PPK-W and PPK-B, respectively. After the quality filtering step, nearly 55% and 29% of the total genotypes of PPK-B and PPK-W samples were retained (**Error! Reference source not found.**).

For the identification of the cause for the melanism we analyzed the ten genes related to the melanin synthesis pathway as well as genes related to phenotypic variation in the coat (Agouti Signaling Protein coding gene.(*ASIP)* (Gene ID 109272540), Melanocortin-1 receptor (*MC1R)* Gene ID 121942601) tyrosinase-related protein 1 coding gene (*TYRP1)* (Gene ID 109247101), Dopachrome Tautomerase Protein coding gene *DCT* (Gene ID 109264168), OCA2 melanosomal transmembrane protein coding gene.*OCA2* (Gene ID 109258721), Phospholipase C Beta 4 protein coding gene*PLCB4* (Gene ID 109274721), Premelanosome Protein coding gene, *PMEL* (Gene ID 109270307), and egl-9 family hypoxia inducible factor 1*EGLN1* (Gene ID 109248931) gene). Out of the analyzed genes, only *ASIP* showed a distinctive variation among the regular leopards and the black leopards. We aligned our two *ASIP* coding sequences (PPK-W, PPK-B) to those previously for available leopard data in NCBI. The Alignment consisted of 408 bp (136 codons) that exhibited heterogenous patterns of variation (Figure 6a). The coding region of *ASIP* gene is a highly conserved within Feliade species, which most of the individuals exhibiting an identical sequences with few exceptions. A Single nucleotide polymorphism (SNP) was identified in exon 4 in the Sri Lankan black leopard. The mutant allele derives a non-synonymous substitution at nucleotide 353 (C353A) predicted to cause a cycteine-phenylalanine substitution at codon 113 (Figure 6b).

**Figure 6:**
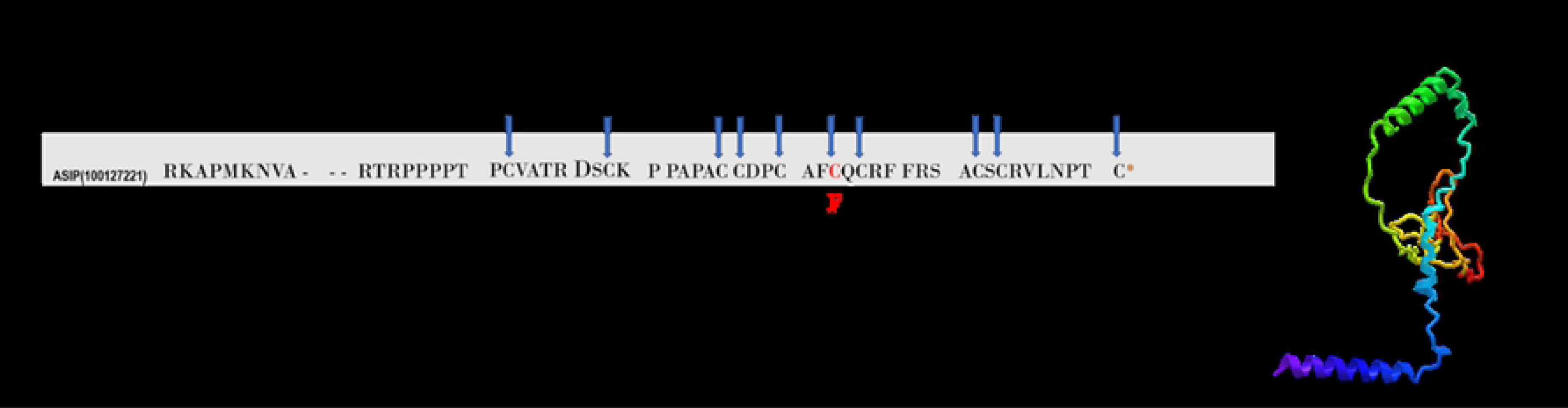
Amino acid alignment of ASIP, including the novel *P.p. kotiya* sequences. (a) Codon Variation in the ***ASIP*** gene Associated with Melanism in the Sri Lankan leopard *P..p. kotiya* Amino acid alignment of *ASIP* Exon 4, including the novel **PPK-W** *P.P. kotiya*- wild and **PPK-B** *P.p kotiya-* black. **ERR5671301** Regular-type *P.p. kotiya* fossil **JX845175** *Panther pardus*. Dots indicate identity to the top sequence; amino acid positions are shown at the end of each line. Dashes represent insertion/deletion (indel) variants. Numbers 1–10 refer to the 10 conserved cysteine residues present in the C-terminal domain. The non-synonymous mutation in melanistic **PPK-B** is indicated in bold and red. (b) Protein structure of ASIP gene in the Sri Lankan black leopard

## DISCUSSION

Though it is illegal in Sri Lanka, not very rare to find traps in the wilderness to kill leopards. However, it is extremely rare to see a trapped black leopard alive and suffering. One such occasion reported on 20th July 2020, marked the third reporting of dead black leopards. While few reports cite black leopards in Sri Lanka, especially in humid upcountry forests, no formal studies so far. There are two different hypotheses among environmental groups, whether it is a different subspecies or of the same subspecies with a black coat. Our study was based on 100x coverage Illumina data generated from the heart blood of a dead black leopard collected during the postmortem and morphological data collected from the same animal. In addition, we generated 30x coverage Illumina data from a regular coat colour animal and morphological data from the same and a few others. While we could only include one animal from each phenotype in the molecular study, we assessed both the mitogenome and the nuclear genome.

The complete mitochondrial DNA sequences provide evidence for molecular dynamics, comparative genetics, and species evolution (55). We chose mtDNA due to its small genome size, maternal inheritance, extremely low probability of paternal leakage, higher mutations rate than nuclear DNA, and change mainly through mutation. We *de novo* assembled the mitogenomes of *P. p. kotiya* (16,847 bp) and black leopard, filling a clear gap in understanding the diversity of *Sri Lankan subspecies.* Although the divergence study highlights the bifurcation between African and Asian leopards and does not confirm the exact origin of *Panthera pardus,* the combined fossil, and genetic evidence together have supported that Africa is the place of origin (1, 5). All the samples in this analysis clustered within the Asian Leopard clade. Our analysis agrees the previous work that Sri Lankan leopards originated from the migration of the Indian subcontinent (3). The estimated coalescence time for Sri Lankan leopards is relatively recent. However, there is no distinct difference in the divergence between the regular and black leopards in Sri Lanka.

Our analysis is consistent with that of Miththapala in 1991 (56), where the Sri Lankan leopards consist of a low heterozygosity and polymorphism. Previous studies found low haplotype diversity in leopards living in West Asian mountains, such as Pakistan, Iran, and Turkmenistan, based on the *NADH-5* gene (57). Our analysis with the full mitogenomes (excluding the D-loop region) confirmed the low variability in mitogenomic sequences of Sri Lankan leopards, compared to Indian leopards. We excluded the D-loop region because of its high complexity and the occurrence of high repeat regions. The nucleotide variation between two samples is relatively low, having only six polymorphic nucleotides in the complete mitogenomic region considered. In particular, we selected the *NADH-5* gene previously used in subspecies delimitation of carnivores, showing a higher mutation rate for this group. All the phylogenetic trees showed that the Sri Lankan black leopard was tied to the same lineage of the Sri Lankan normal leopard. The Sri Lankan regular leopard and black leopard clustered together. Therefore, we propose that the Sri Lankan regular and black leopards belong to the same subspecies.

However, the haplotype network analysis conducted with current sequences and previously purchased highlighted the presence of two different haplotypes with a single nucleotide variation. The regular coat colour (BBK-W) and black leopard (BBK-B) belong to the same haplotype. The previously recorded sequences, (AY035268.1 and AY035269.1) were grouped into a separate haplotype. The genetic variation of the Sri Lankan leopard samples did not support the existence of a another subspecies in Sri Lankan, whether existing or novel. On the other hand, subspecies of *P.p. kotiya* may have a low genetic variation, suggested by Uphyrkina and colleagues grouped the *kotiya* leopard into three haplotypes (1). Nevertheless, our data suggest that the Sri Lankan black leopard is evolutionarily close to the common coat colour leopard in Sri Lanka. Including more samples in the analysis is needed to confirm this hypothesis. It will give a clear idea about the intraspecies diversity among Sri Lankan leopard populations.

We tested the hypothesis of whether the colouration is due to a mutation or mutations in the melanin synthesis pathway. Previously proposed that the Leopard colouration possesses adaptive relevance, performing various roles in behavioral and ecological processes (37, 58). The density of melanin and the distribution of melanin types on individual hairs determine the coat colour of mammals including leopards (37). Biological factors such as camouflage, thermoregulation, sexual selection, reproductive success, and parasite resistance could also be the reasons associated with the melanism in leopards (59, 60). The occurrence of melanism in leopards has been documented in 13 out of its 37 species (41, 61). The darker coat colour is considered adaptive to wetter areas with dense vegetation (34, 62). Even in Sri Lanka, black leopards have mainly been reported in tropical and humid environments. To clarify the exact reason behind the melanism, we analyzed coding sequences of the genes related to the melanin synthesis pathway and receptor genes.

In leopards, extreme phenotypic changes in the coat colour patterns are associated with single mutations of *MC1R* and *ASIP* genes (60). The *MC1R* is a single-exon gene, and mutations change its amino acids that alter the receptor’s affinity for binding with ligands and G-protein. Eumelanin, the dark pigment, production driven by the *MC1R* gene is activated by the binding of Alpha Melanocyte Stimulating Hormone (α-MSH) (63). The *ASIP* gene consists of a peptide of 131 amino acids with a conserved cystein-rich region and a putative signal peptide at the N-terminus (64). The cysteine-rich domain-containing ten (10) cysteine residues are conserved across taxa (64, 65) and are responsible for its biological activity as an inverse agonist of *MC1R* (39, 66). The *ASIP* C-terminal loop, a six amino acid segment closed by a final disulfide bond, is essential for high-affinity *MC1R* binding and inhibits the activation of *MC1R* for the production of Eumelanin and leads to produce pheomelanin (67). Therefore, the gain of function in *MC1R* or loss of function in *ASIP* induces melanism.

Our results suggest that SNP identified in the *ASIP* case the melanism in *P.p kotiya. Because* of the mutation, the 6th C-residue in the C-terminal loop is substituted by Adenine. It may result in complete loss of function of the *ASIP,* inhibiting the binding of *ASIP* to the *MC1R*. Therefore, eumelanin production does not inhibit, resulting in a black-coated leopard. Additional replicated functional studies are thus required to assess in detail the pleiotropic effect in the loci. However, in the Sri Lankan leopard, the black rosettes are still visible despite much-darkened background colouration. It is distinct in the cross-section that the beginning of the coat shaft has a cream colouration (Fig 1b). It suggests that it is a mutation of the regular coat of the Sri Lankan leopard, and rosettes are still darker than the darker fur in the background. The functional assays will directly establish the biological effects of these variants. Though several other mutations in the same gene are reported in several other instances, a nonsense mutation in exon 4 (C333A) and a non-synonymous substitution (C384G) (40). However, the same mutation (C353A) is not reported before. Therefore, this can be considered a novel mutation identified in the ASIP responsible for polymorphism in Felidae family.

Consequently, our study suggests that the black leopards in Sri Lanka do not appear to have any significant genetic speciation or endemic heredity. Nevertheless, they play role in ecological overlap within nature and provide suitable conditions for a high level of gene flow between them. In Sri Lanka, melanism is recorded at low frequencies, in contrast to other countries. According to Kingsley (60), the *ASIP*-induced melanism is recessive. As a result, it would take a longer time to rise in frequency when favorable and may linger in the population for a longer period when negatively selected. On the other hand, the *ASIP-induced* melanism can reach high frequency in some populations, suggesting that the trait may be adaptive or at least neutral. Therefore, there is a high probability of reoccurrence of black leopards in Sri Lanka if the habitat is favorable. One of the main constraints in the conservation of leopards in Sri Lanka is lack of detailed information, both on black leopards and regular coated leopards in highly fragmented habitats. Our findings show the importance of cutting-edge technologies for a deep understanding of the germplasm for comprehensive conservation decisions. Including more samples will provide a better resolution.

## CONCLUSION

Accordingly, Sri Lankan black leopard appears to be from the same lineage as the Sri Lankan regular leopards and has acquired the colour just because of a mutation. This may be mainly because of the environmental adaptation. The Sri Lankan black leopards have a different mutation compared with the other recorded *Panther pardus* black leopards. The SNP detected in *Panthera pardus kotiya* causes an amino acid change (C353A) that induce a loss of faction in the *ASIP* gene. Therefore, our study reveals an additional case of species-specific mutation caused melanism in the Feliedae family. As such, *ASIP* mutation may play an important role in naturally occurring colouration polymorphism. Since this mutation may be due to the camouflage, it has a high probability of reoccurrence and inheritance to the next generations. Moreover, our assembled genome could be improved further and annotated using advanced sequencing approaches like HiC and transcriptome assembly.

## Declarations

### Ethical Approval

The study was conducted under the research permit number: WL/3/2/2017/1 issued by the Research Committee of the Department of Wildlife Conservation, Sri Lanka. All the methods were carried out in accordance with relevant guidelines and regulations for animals. The DNA samples for sequencing were exported under CITES (Convention on International Trade in Endangered Species of Wild Fauna and Flora) permit (Security Stamp 1888968) obtained from the DWC.

### Consent for publication

Not applicable

### Availability of data and materials

The regular and black leopard whole genome project has been deposited at DDBJ/EMBL/GenBank. The scripts and codes used in this study is available in https://github.com/AgBC-UoP/sri-lankan-leopard

### Authors’ Contributions

**M.G. Chandana Sooriyabandara** Funding acquisition, Investigation, Sample submission

**Asitha U. Bandaranayake** Conceptualization, Data curation, Formal analysis, Investigation, Methodology, Software, Validation, Visualization

**Shyaman M. Jayasundara** Conceptualization, Data curation, Formal analysis, Investigation, Methodology, Validation, Software, Visualization, writing-original draft, Writing –editing

**H.A.Bhagya M. Hathurusinghe** Conceptualization, Data curation, Formal analysis, Investigation, Methodology, Validation, Visualization, writing-original draft, Writing –editing

**M.S.L. Ranjan P. Marasighe** Funding acquisition, Investigation, Sample submission

**G.A.Tharaka Prasad** Sample submission

**V.P.Malaka K.Abeywardana** Sample submission, Methodology

**M.A. Pinidiya** Sample submission, Methodology

**R.M. Reunka Nilanthi** Conceptualization, Investigation, Sample submission

**Pradeepa C. G. Bandaranayake** Conceptualization, Data curation, Formal analysis, Investigation, Methodology, Project administration, Resources, Software, Supervision, Validation, Visualization, writing – original draft, Writing – review & editing

## Acknowledgements

We especially thank the field staff of the DWC for the collection of samples and the staff of Agricultural Biotechnology Centre, University of Peradeniya, Sri Lanka especially, Dr Bhagya Chandrasekara for their immense support throughout the Project.

## Additional Files

Additional File 1: Table S1: Genetic contents in the mitochondria genome of *P. pardus kotiya. Docx*

Additional File 2: Figure S1: GC content test for the reverse reads of *P. p. kotiya* library.Docx

Additional File 3: Table S2: Morphometric measurements of PPK-W and PPK-B leopards.Docx

